# Large-scale Manual Curation and Harmonization of Metadata from Metagenomic and Cancer Genomic Repositories: Challenges and Solutions

**DOI:** 10.1101/2025.11.26.689816

**Authors:** Kaelyn Long, Kai Gravel-Pucillo, Levi Waldron, Sean Davis, Sehyun Oh

**Affiliations:** Institute for Implementation Science in Population Health, City University of New York School of Public Health, New York, NY; Department of Epidemiology and Biostatistics, City University of New York School of Public Health, New York, NY; Departments of Biomedical Informatics and Medicine, University of Colorado Anschutz School of Medicine, Denver, CO

**Keywords:** OmicsMLRepo, FAIR Principles, Clinical Data, Metadata Harmonization, AI/ML-ready, cBioPortal, curatedMetagenomicData

## Abstract

Public omics repositories contain vast amounts of valuable data, but their metadata suffers from extreme heterogeneity, unstandardized terminologies, and quality issues that severely limit data reusability and cross-study integration. While prospective metadata standards exist, the majority of published omics data remain in non-standardized formats requiring retrospective curation. We performed comprehensive manual curation and harmonization of clinical metadata from 212,027 samples across 468 studies in two major repositories: *curatedMetagenomicData* (93 studies, 22,588 samples) and cBioPortal (375 studies, 189,438 samples). Through systematic ontology mapping, we consolidated redundant, dispersed information into much fewer harmonized columns, reduced unique values, and increased the completeness of major attributes. This curation process revealed common metadata quality issues, including typos, inconsistent terminologies, misplaced values, conflicting annotations, and inappropriately merged information across attributes. We document the challenges, decisions, and solutions encountered during large-scale metadata harmonization across two distinct omics domains. The harmonized metadata, accessible through the *OmicsMLRepoR* Bioconductor package, enables repository-wide queries and cross-study analyses previously challenging with heterogeneous metadata. Our experience provides practical guidance for similar curation efforts and demonstrates the value of investing in retrospective metadata improvement for existing public omics resources.

## Introduction

The exponentially growing volume of public omics data presents opportunities for cross-study analysis, thereby improving statistical power and enhancing generalizability (1). Several examples have demonstrated significant advantages in understanding diseases through cross-study analysis, including the identification of disease signatures(2), enriched pathways(3), and potential treatments(4,5). This potential requires standardized metadata that enable data identification, integration, and interpretation; however, their innate complexity and heterogeneity pose a significant challenge to standardization and validation (6–8). Despite numerous calls for improved metadata standards and several proposed frameworks (9–11), the reality is that most published omics data still exist with heterogeneous, unstandardized metadata that severely limits its usability.

Several challenges exist in developing metadata that complies with the FAIR (Findable, Accessible, Interoperable, Reusable) and TRUST (Transparency, Responsibility, User focus, Sustainability, and Technology) principles (9). First, there are too many data standards(10). Some existing data standards impose overly granular specification requirements that exceed practical implementation capacities(11). Additionally, the metadata augmentation efforts often focus on developing new data standards(12,13) rather than implementing existing ones. Second, biological data collection and management commonly involve unstandardized values, such as curator-defined abbreviations and drug brand names. The same or similar concepts are frequently categorized or labeled differently across studies. Ontologies provide a solution by offering standardized vocabularies and identifiers for domain-specific concepts, facilitating data organization, integration, exchange, and reuse in machine-readable formats. However, ontologies are not widely used at the biomedical data collection stage due to several practical barriers, including their complexity and training requirements, time constraints in clinical settings, and software limitations such as EHR tools that lack ontology integration. Third, most biomedical repositories lack quality and sanity checks at the value level. Common human errors, such as typos or values under the wrong attribute, can be systematically recognized and corrected. However, no convenient utility exists for performing those checks, particularly when varying patterns exist across studies.

Several standardization frameworks have been developed to address metadata heterogeneity. The Observational Medical Outcomes Partnership (OMOP) Common Data Model(14,15) provides a comprehensive framework for standardizing observational health data, particularly electronic health records. Health Level Seven Fast Healthcare Interoperability Resources (HL7 FHIR)(16) offers flexible, web-based standards for health information exchange. While these standardization frameworks have advanced metadata quality in their respective domains, none adequately addresses the critical challenge of retrospective harmonization across heterogeneous omics research repositories. OMOP CDM and HL7 FHIR require prospective adoption and infrastructure changes that are impractical for the vast quantities of already-published research data. At the same time, their clinical focus makes them less suited for the diverse experimental designs and biological contexts inherent to omics studies. The ISA framework and domain-specific standards (e.g., MIAME(17), MINSEQE (https://www.fged.org/projects/minseqe/), MIxS(18)) excel at guiding prospective metadata collection but offer limited guidance for reconciling the inconsistent terminologies, varying granularities, and conflicting annotations that characterize legacy datasets. Previous retrospective harmonization efforts have remained confined to small numbers of carefully selected cohorts within single research domains, typically harmonizing 3-10 studies with relatively homogeneous data structures(19,20). No existing framework provides a systematic methodology for retrospectively harmonizing metadata across hundreds of independently conducted studies spanning multiple omics types, each with distinct data models, terminologies, and curation practices. This gap is particularly problematic given that the majority of valuable public omics data already exists in non-standardized formats, and achieving consensus for prospective standards across the diverse omics research community remains challenging. There is an urgent need for a lightweight, practical approach specifically designed to harmonize heterogeneous legacy metadata at repository scale without requiring retroactive infrastructure changes or forcing artificial conformity to overly rigid prospective schemas.

To address this gap, we introduce the OmicsMLRepo project - a metadata harmonization initiative focusing on the practical and retrospective harmonization of already-collected research data across multiple omics domains (**Table 1**).

**Table 1.** Comparison between OmicsMLRepo and existing standardization frameworks. It highlights key differences in application scope and requirements.

| Framework | Prospective/<br>Retrospective | Scale | Domain | Infrastructure<br>Required | Primary Use<br>Case |
| --- | --- | --- | --- | --- | --- |
| OMOP CDM | Prospective | Large | Clinical | High | EHR<br>standardization |
| HL7 FHIR | Prospective | Any | Clinical | Medium-High | Health data<br>exchange |
| ISA | Prospective | Study-level | Lab/<br>Experimental | Medium | Metadata<br>collection |
| OmicsMLRepo | Retrospective | Repository-<br>level | Multi-omics<br>research | Low | Legacy data<br>harmonization |

Here, we report our experience performing large-scale manual curation and harmonization of clinical metadata from two major omics repositories: *curatedMetagenomicData* (cMD)(21) and *cBioPortalData*(*22*) Bioconductor packages. The cMD package contains 22,588 whole metagenomic shotgun sequencing datasets from 93 studies, all uniformly processed and accompanied by manually curated metadata (21). While cMD metadata were curated by experienced curators, it has not been harmonized across the studies and are still plagued by low-quality issues, such as typos and redundancy. The *cBioPortalData* package provides access to data from the cBio Cancer Genomics Portal (cBioPortal)(23–25). cBioPortal is a free, open-source platform for exploring and analyzing cancer genomics data, providing large-scale cancer genomics datasets that include molecular and clinical metadata from various studies and sources. However, cBioPortal’s clinical metadata is extremely heterogeneous and lacks standardization. Different studies employ distinct and often arbitrary terms and abbreviations for identical entities, making it challenging to identify samples across studies based on clinical information.

The OmicsMLRepo project augments metadata by performing data validation and correction, identifying and consolidating dispersed information into parsimonious representations, and mapping values to existing ontologies. We demonstrated the improved usability of the harmonized metadata through examples of repository-wide summary statistics. We developed the *OmicsMLRepoR* Bioconductor package for easy access and manipulation of harmonized metadata, while redistributing harmonized metadata through their original repositories as well to minimize data replication. The OmicsMLRepo project will enable researchers to effectively find and utilize relevant datasets, reduce redundant data generation, facilitate cross-study analyses, improve AI/ML-readiness of public omics data, advance research reproducibility, and accelerate discoveries.

## Methods

### Datasets

We harmonized metadata from the *curatedMetagenomicData* (cMD, version 3.8.0) and *cBioPortalData* Bioconductor packages. For *cBioPortalData*, we captured sample-level clinical metadata on May 8, 2023, and selected core attributes for harmonization based on their completeness, prevalence across data resources, and clinical relevance. All the cMD attributes were harmonized. To minimize data duplication and changes in the existing workflow, our harmonized metadata is implemented back to the source packages.

### Exploratory data analysis and schema development

Before establishing our harmonization schema, we conducted comprehensive exploratory data analysis (EDA) to understand the heterogeneous structure of metadata across studies. This process involved examining both attribute names and their corresponding values, as datasets exhibited significant structural differences in how they organized conceptually similar information. For example, one study might consolidate all treatment information into a single column containing medication name, type, dosage, and administration frequency as free text, while another study might decompose the same information into separate binary attributes for each specific treatment. We systematically identified and grouped related information scattered across different attributes, which enabled us to interpret individual values within their proper context, apply consensus-based harmonization strategies, and minimize redundancy in the final harmonized dataset. This data-driven approach, in contrast to imposing predetermined schemas, allowed us to develop a data model that accurately represents real-world data characteristics and facilitates integration of diverse data sources.

### Extract, Transform, and Load (ETL)

During the data extraction step, we manually reviewed and identified information dispersed and redundant across multiple attributes, collecting all unique values from them. These values were manually curated to ontology terms, and the resulting per-attribute ’*original value: ontology term*’ map was used for value transformation. For numeric attributes, we separated the numeric and unit components.

The harmonization process involved systematic data cleaning and ontology mapping procedures:

● Data cleaning:

- Multiple values assigned to a single cell were delimited using semicolons unless otherwise specified
- Binary columns (e.g., presence/absence) were converted to descriptive values
- Empty strings and non-interpretable values were assigned *NA*
- When multiple values at different resolutions were available for a given term, we consolidated to the most specific information
● Ontology mapping:

- Full names were identified for all abbreviations
- Unique values were mapped to appropriate ontology terms(26)
- Original values were updated using manually created ’*original value: ontology term*’ pairs

We also revised inaccurate and conflicting information through cross-comparison and literature reviews. When there were multiple ontologies for a single term, we prioritized them based on two criteria: 1) one leading to fewer ontologies per attribute and 2) one that provides better interoperability with other resources.

For each metadata attribute, all unique original values were manually mapped to curated terms. These curation maps include four fields: original_value, curated_ontology_term, curated_ontology_term_id, and curated_ontology_term_db. Exceptions included partially different column names (e.g., curated_antibiotics), additional columns when needed (e.g., curated_age_min and curated_age_max for age group curation), and newly created columns (e.g., curated_target_condition). The consolidation process and resulting data model are summarized as merging schema tables (**Supplementary Tables 1-2**) and data dictionaries (**Supplementary Tables 3-5**), respectively.

### Conflict resolution

Harmonization of metadata from multiple sources inevitably revealed conflicting annotations for the same samples. We developed a systematic approach to resolve these conflicts based on the nature and cause of discordance. First, we distinguished between systematic and random errors, as this determines the appropriate correction strategy. For example, during harmonization of the sex attribute in *cBioPortalData*, we found that 303 out of 337 discordances originated from a single study (*luad_msk_npjpo_2021*), which we confirmed as a systematic error at the original data recording step. Such systematic errors were corrected at the source level when possible. Second, we applied logical consistency checking through cross-attribute validation based on established medical knowledge. For instance, combinations such as ’prostate cancer’ with ’female’ sex represent logical impossibilities that flagged errors requiring correction. Third, for attributes that remain constant over time (e.g., sex assigned at birth, genetic ancestry), we applied consensus-based harmonization using a majority rule. When multiple samples from the same patient contained conflicting values for such attributes, we selected the value appearing most frequently. If no majority existed, we replaced values with *NA*. This approach enabled us to infer sex for 422 additional samples and correct 39 incorrect values in the cBioPortal metadata. We carefully limited this consensus-based strategy to attributes that cannot legitimately change over time, avoiding its application to dynamic clinical variables such as disease stage or treatment status.

### Ontology integration and dynamic value validation

We leveraged ontologies not only for standardizing values but also for systematic value validation. For each harmonized attribute, we defined one or more “dynamic enumeration nodes” (*dynamic enums*). Dynamic enum nodes are selected among the shared ancestors from relevant ontologies (i.e., already captured values). All the descendant terms from dynamic enums automatically constitute the allowed values for that attribute. For example, we designated ’ancestry category’ (HANCESTRO:0004) as a dynamic enumeration node for the ancestry attribute, making all eight child terms and their combinations valid values. When choosing dynamic enums, we carefully evaluated the trade-off between creating categories that are too broad to be useful versus too narrow to accommodate diverse data sources. We addressed the challenge of rapidly emerging concepts not yet incorporated into established ontologies by creating corresponding ’*_details’ attributes to preserve granular information that lacks standardized ontology terms.

We can systematically identify data entry errors by validating values against a pool of allowed values (i.e., descendants of dynamic enums); if values do not belong to the pool, they are likely wrong or miscategorized values. We also validate manual curation results by looking up ontology terms using curated ontology IDs (via the *rols* R package(27)) and comparing them with the curated ontology terms. Updates were frequently made during the sanity check, including correcting typos, incorrect interpretations of abbreviations, incorrect ontology IDs, and mismatches between ontology terms and IDs. This ‘round-trip validation’, as well as the dynamic enum-based validation, enabled us to identify and correct inconsistencies that would have otherwise compromised data quality.

### Flexible attribute structures for heterogeneous data

The exploratory data analysis revealed that rigid single-valued schemas could not accommodate the structural diversity across studies. We therefore implemented three attribute types based on their value structure: single-valued attributes (one value per attribute), multi-valued attributes (multiple values per attribute), and composite attributes (multiple related features under a single generic attribute). Multi-valued attributes allow multiple standardized values to be assigned to a single attribute for a single sample (e.g., a patient receiving multiple concurrent treatments). Values are separated by semicolons as delimiters. This design preserves the complete information without requiring separate rows for each value, which would complicate sample-level analyses. Composite attributes enable logical grouping of related features under a single generic attribute. This structure improves conceptual clarity and reduces visual complexity in data tables, enables users to quickly understand the extent of available data, provides flexible representation that can accommodate different numbers of measurements per sample, and allows easy addition of new measurements without restructuring the entire database.

To address granularity mismatches across data sources, we created paired attributes: standard/main attributes containing harmonized terms at an appropriate level for cross-study comparisons, and corresponding ’*_details’ attributes preserving more specific information. For instance, the ancestry attribute contains eight standardized child terms from HANCESTRO:0004, while ancestry_details preserves additional granular information mapped to extended ontology terms. This dual-level approach eliminates the traditional trade-off between preserving granular detail and achieving broad compatibility, ensuring that no meaningful information is discarded during harmonization.

### Naming schema and data provenance

To maintain transparency and traceability, harmonized metadata tables include: 1) columns with ’original_’ or ’curated_’ prefixes (in the data provenance version), 2) a ’last_modified’ timestamp, and 3) a ’curation_id’, which is a concatenation of other ID columns to create a unique identifier per sample (study_name:participant_id for cMD; study_name:participant_id:sample_id for *cBioPortalData*). The naming schema for data provenance follows this structure:

- curated_{attribute_name} (e.g., curated_age)

- curated_{attribute_name}_{options} (e.g., curated_age_unit)

- curated_{attribute_name}_ontology_term_id

(e.g., curated_age_ontology_term_id)

- curated_{attribute_name}_source (e.g., curated_age_source): the original attribute name from which the value was captured

- original_{attribute_name}_value (e.g., original_age_value)

### Data dictionary

The data dictionary describes the major characteristics of the curated fields (**Supplementary Tables 3-5**). We maintain two formats for the same version: one for curators during the data curation stage and another for the release version presented to end-users. The main difference is compactness of display—for curation versions, we split composite attributes to facilitate detailed editing, while release versions present them in their grouped form.

### Merging schema

A merging schema table summarizes how original fields/attributes were harmonized into curated fields/attributes and records their completeness and variability before (’original_*’) and after (’*curated_’) harmonization. For cMD, a single merging schema table contains six columns: original_field, original_field_completeness, original_field_unique_values, curated_field, curated_field_completeness, and curated_field_unique_values. For *cBioPortalData*, the schema is structured similarly but accounts for the many-to-many mapping relationships between original and curated attributes.

### *OmicsMLRepoR* package implementation

The *OmicsMLRepoR* Bioconductor package enables users to navigate harmonized metadata through ontological relationships in a robust and user-friendly manner. The main function, tree_filter, performs searches that include not only the queried terms but also their synonyms and all descendant terms, leveraging the ontology hierarchies we established during harmonization.

To address use cases where users need subsets of multi-valued and composite attributes, *OmicsMLRepoR* provides metadata reshaping functions—spreadMetaTb and gatherMetaTb. These functions spread or gather the metadata table anchored around selected attributes, transforming between wide and long formats to enhance downstream analysis capabilities. This design allows users to maintain both sample-focused and attribute-focused analytical workflows from the same underlying harmonized data structure.

## Results

We harmonized clinical metadata from 212,027 samples across 468 studies available through the cMD and cBioPortalData packages (**Figure 1a**). The quality of harmonized metadata was enhanced through de-replication, ontology incorporation, and reconciliation. We assessed the improved metadata quality and the impact of harmonization using four criteria (**Figure 1b**): 1) attribute standardization, representing the consolidation of variables and measured as the number of original attributes merged into each new curated attribute (compression); 2-1) value standardization, measured as the number of unique values per attribute (consolidation); 2-2) value correction, represented as the proportion of values updated during curation (correction rate); and 3) the combined impact of attribute and value standardizations, demonstrated through the completeness of the attributes (completeness), represented as the proportion of the non-*NA* values per attribute.

**Figure 1.**
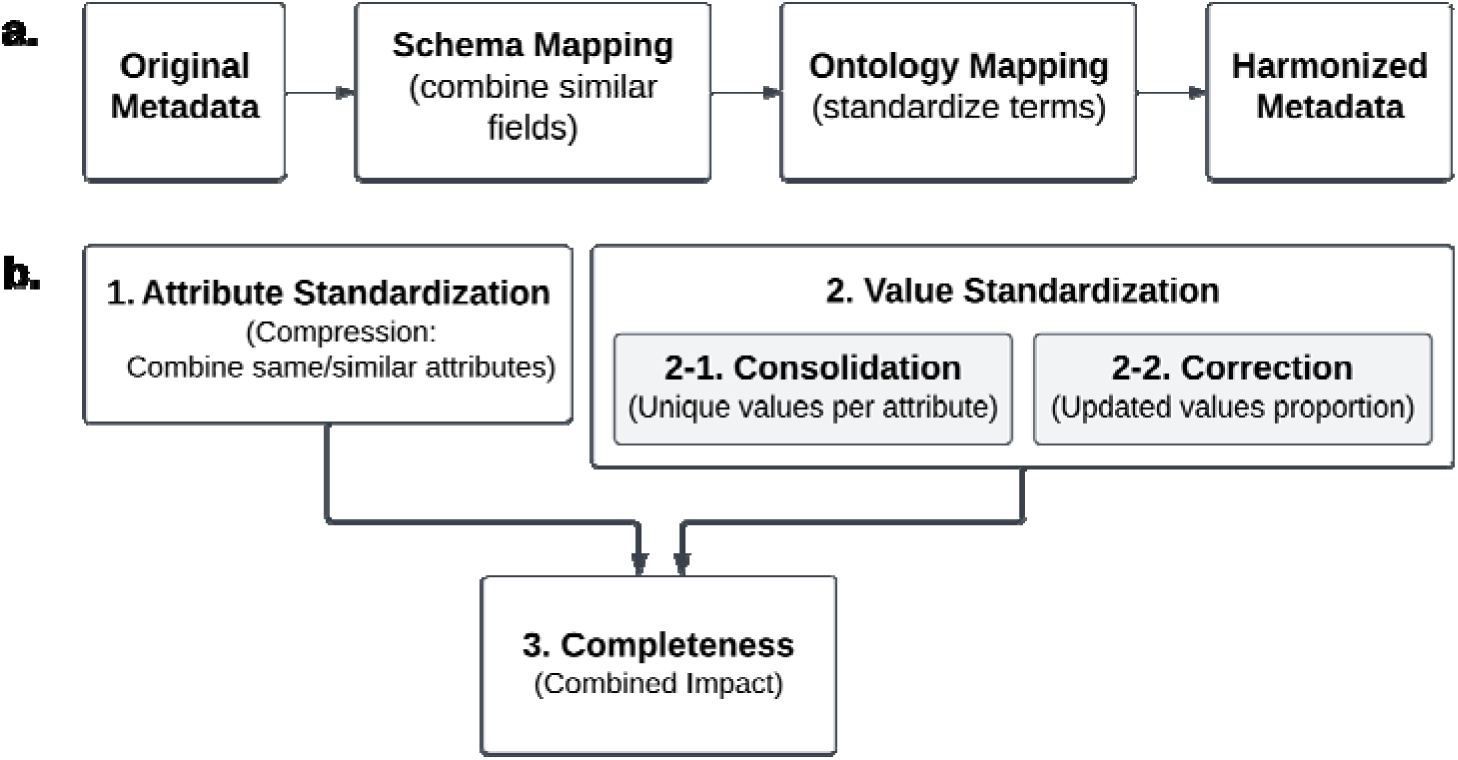
Overview of the harmonization process and quality metrics. **(a)** Simplified workflow showing the major steps in metadata harmonization. The process transforms original heterogeneous metadata into harmonized, ontology-mapped values. **(b)** Four metrics used to assess improved metadata quality.

### Augmented metadata quality for *cMD*

After harmonization, 142 original attributes were compressed into 66 attributes (**Figure 2a**). The biomarker exhibited the most data compression, as 38 original attributes for different biomarkers were merged into a single composite attribute, *‘biomarker’*. The harmonized metadata improves the completeness of individual attributes (**Figure 2b**). For example, the curated *‘hla’* column reached ∼5% complete, while each of the five original columns about *hla* genotypes (*HLA*, *hla_dqa11*, *hla_dqa12*, *hla_drb11*, and *hla_drb12*) was less than 2% complete. We observed a very high correction rate in cMD harmonization because most of the categorical variables used non-standardized terms. For example, over 97% of values in the original *disease* and *treatment* columns were updated to ontology terms in their curated columns, which improves interpretability, findability, and interoperability (**Figure 2c and Table 2**). The cMD metadata usability was further improved by reducing redundancy, correcting mismatches, reorganizing values under proper attributes, and fixing typos. Our harmonization process also untangles intertwined and buried information. For example, the original cMD metadata had two poorly defined attributes (*study_condition* and *disease*) that contained mixed information. The harmonized schema separates these into three clearly defined attributes (*control*, *target_condnition*, and *disease*), enabling users to identify unconventional situations such as non-healthy participants serving as controls or differentiate two participants with the same conditions (e.g., having both type 2 diabetes and adenoma) subjected to different studies (e.g., studies targeting type 2 diabetes versus adenoma), which will require distinct interpretations.

**Figure 2.**
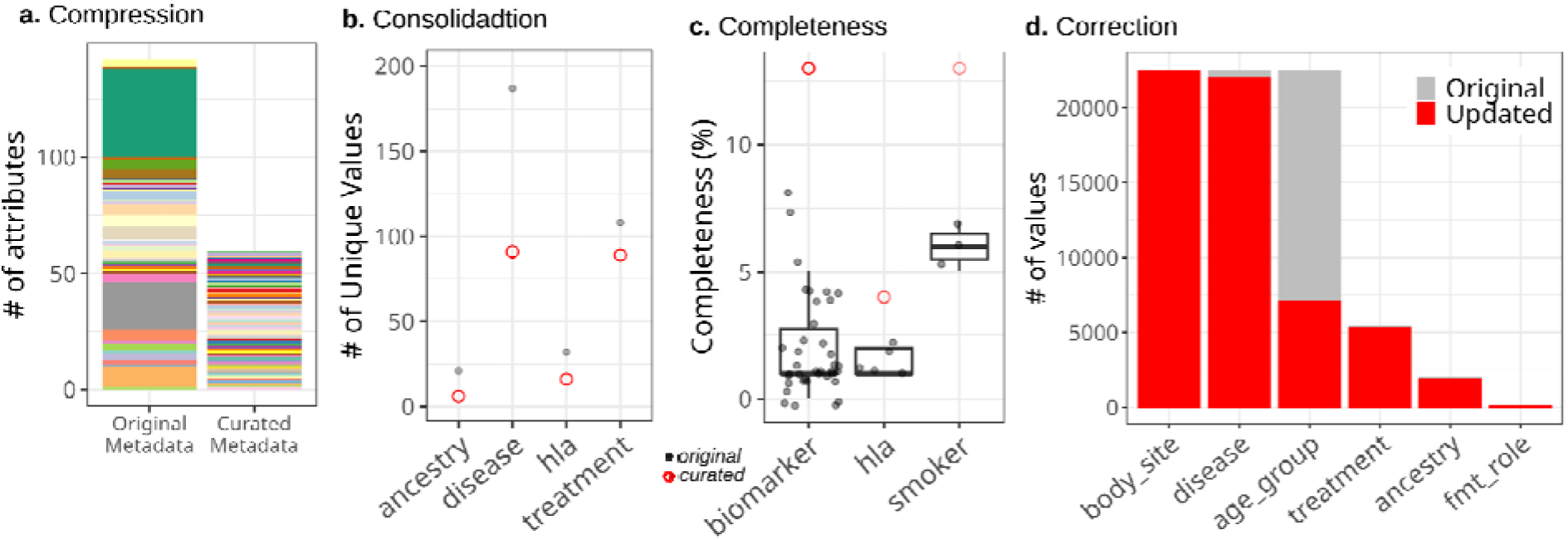
Improved metadata quality for *cMD*. **(a)** The color schema used in the left bar (’Original Metadata’) represents the original attributes and the corresponding curated attributes. **(b-d)** The x-axis shows curated attributes. The grey points in panels b and c represent the original attributes, and the red circles indicate the curated attributes. **(d)** The number of values that were kept in their original form (grey) and updated during harmonization (red) per attribute was plotted.

**Table 2.** Examples of original and curated values. The curated *treatment* column (on the right) contains more detailed and accurate information than the original *treatment* values (on the left). The standardization involves re-recording values (e.g., brand name, such as ‘Lantus Solostar’, which was initially entered separately as ‘lantus’ and ‘solostar’) into a raw generic drug (e.g., ‘Insulin Glargine’). These are examples from the *cMD*.

| Original *treatment* column | Curated *treatment* column |
| --- | --- |
| antilipid;antihta;antidiab;diuranse;ace_inhib;beta_blockers;ca2_cbl;metformin;glp_1;dppiv;insulin;statin;asa;clopidogrel;nitrate;ppi;xantoxinh | Antilipidemic Agent;Antihypertensive Agents;Anti-diabetic Agent;Diuretic;Angiotensin-Converting Enzyme Inhibitors;Beta-Adrenergic Antagonist;Calcium Channel Blocker;Metformin;glucagon-like peptide-1 receptor agonist;Dipeptidyl-Peptidase IV Inhibitors;Insulin;statin;Aspirin;Clopidogrel;Nitrate;Proton Pump Inhibitor;Xanthine Oxidase Inhibitor |
| metformin;sitagliptin;lantus;solostar;novorapid | Metformin;Sitagliptin;Insulin Glargine;Insulin Aspart |

### Augmented metadata for cBioPortal

The snapshot of cBioPortal clinical metadata used for harmonization contained 189,439 samples, ranging from 5 to 25,775 samples per study, from 375 studies, including TCGA, TARGET, and publications from individual labs. Clinical metadata from these samples contained 3,733 attributes, with over 95% attributes having less than 4% data completeness. We harmonized 673 columns containing information on core attributes (i.e., age, body site, sex, disease, treatment, and ancestry) into 30 curated attributes (**Figure 3a**). The maximum data compression was achieved for treatment types, where values across 254 original attributes were condensed into a single curated attribute, *treatment_type* (**Supplementary Table 2**). During the initial round of harmonization, we observed an average of ∼71% reduction in the number of unique values for curated attributes (**Figure 3b**). The curated attributes exhibit significantly improved completeness (**Figure 3c**). About 78% of the original attributes (527 out of 673) were less than 1% complete; however, the harmonized attributes have an average completeness of about 25%, and half of the attributes (15 out of 30) have a completeness above 10%. cBioPortal’s clinical metadata harmonization involved a many-to-many merging process, where one original column frequently contributed to multiple curated columns and vice versa; therefore, our ‘correction rate’ measure was not applicable.

**Figure 3.**
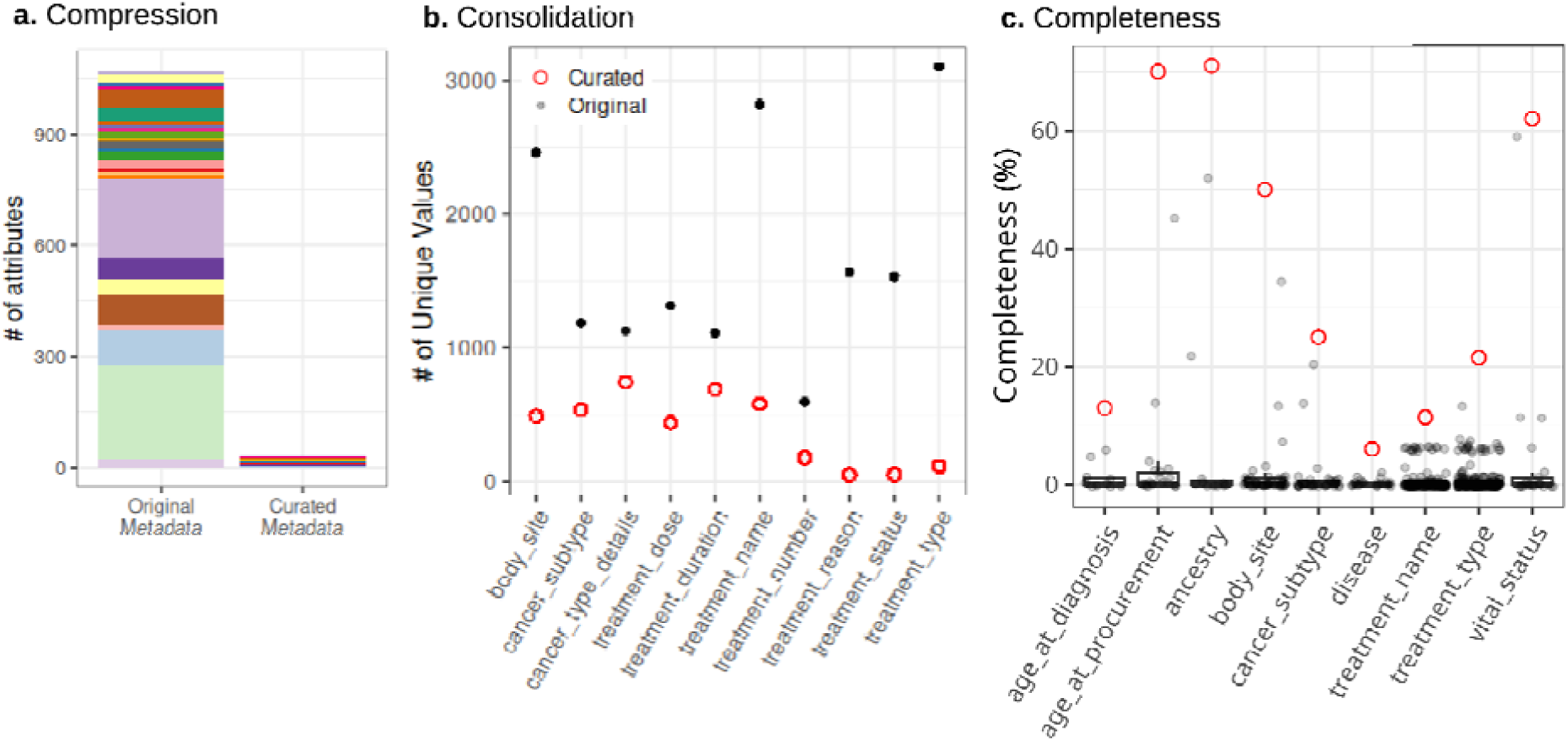
Improved metadata quality for *cBioPortalData*. **(a)** The color schema used in the left bar (’Original Metadata’) represents the original attributes combined into the same curated attribute. The number of source attributes in this plot exceeds 673 because information from some original attributes contributed to multiple curated attributes. **(b)** Top 10 attributes showing the most reduction in the number of unique values, and **(c)** top 10 attributes showing the most increase in their completeness. The x-axis shows the curated attributes. The black and grey points represent the original metadata, and the red circles indicate the curated metadata.

### Analysis of harmonized metadata

Having unified access to summary statistics across all studies in the repository, we can identify patterns and correlations that span multiple datasets. For example, researchers can quickly identify all samples with specific ancestry proportions across different tumor types to study population-specific genomic variations or to identify treatment responses across diverse patient cohorts. Also, we can identify and assemble larger, more diverse cohorts by querying across the entire repository. This is particularly valuable for rare diseases or specific demographic groups where individual studies might have limited sample sizes. A comprehensive view would help researchers understand what data is available repository-wide, preventing duplication of efforts and identifying gaps in the data landscape. This could inform future data collection priorities and help maximize the utility of existing datasets. Additionally, repository-wide summaries would facilitate the identification of inconsistencies in data annotation, missing values, or outliers across studies, thereby making quality validation and standardization easier. The following examples illustrate repository-wide analyses enabled by harmonized metadata.

#### Ancestry

Approximately 71% of cBioPortal samples (n = 134,032) contain information related to ethnicity, race, and ancestry, which were spread across 11 attributes (**Supplementary Table 2**). We harmonized these into two attributes - ancestry and ancestry_details. The ancestry attributes comprise eight child terms from the ‘ancestry category’ (HANCESTRO:0004) ontology term, resulting in 25 unique combinations of ethnic backgrounds (**Figure 4a**). Any values with additional details beyond these eight child terms were mapped to ontologies (HANCESTRO and NCIT) and saved under the ancestry_details column.

**Figure 4.**
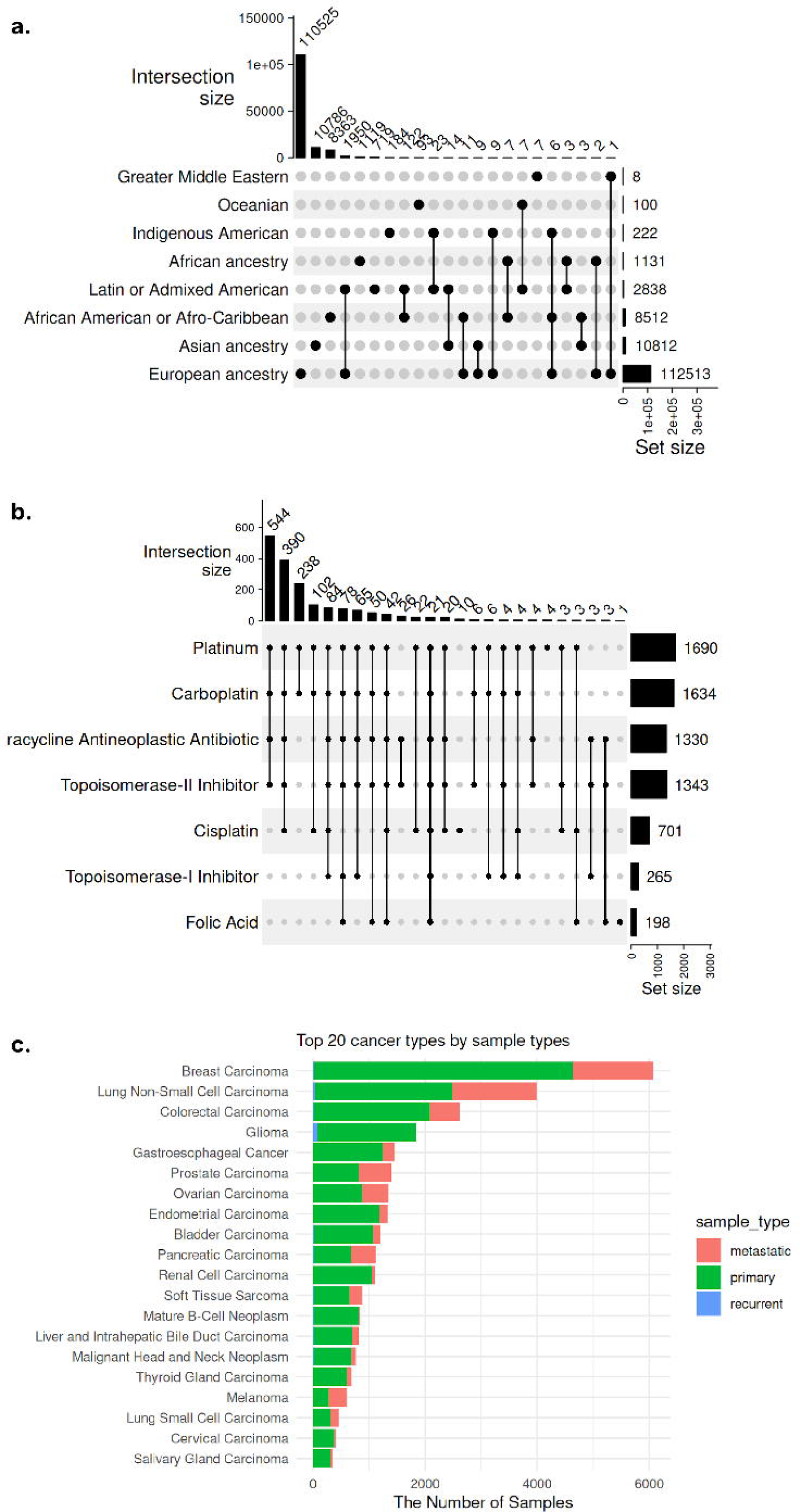
Repository-wide statistics of key attributes. **(a)** A summary of ancestry information of 134,032 samples from 218 studies in the cBioPortal. There are eight child terms of the ‘ancestry category’ (HANCESTRO:0004) ontology term and 25 different combinations of them. (**b**) A summary of treatment names applied to 1,817 samples from 4 independent ovarian cancer studies. **(c)** Available sample types for each cancer type.

#### Treatment Names

Using the harmonized cBioPortal metadata, we can access samples across studies from a single query. For example, we searched all treatment names to which ovarian cancer samples were exposed and found 1,817 ovarian carcinoma samples from 4 studies that contained treatment name information. Platinum is most frequently used in combination with Carboplatin, Tetracycline Neoplastic Antibiotic, and Topoisomerase-II inhibitor (**Figure 4b**).

#### Sample and Specimen Types

In the original cBioPortal metadata, sample- and specimen-type information was spread across over fifty attributes. We harmonized the top three most complete original attributes (SAMPLE_TYPE, SAMPLE_CLASS, and SPECIMEN_TYPE) that were redundant and also mixed with other features, such as disease name and treatment type. In the harmonized metadata, the specimen_type attribute includes only standard terms for categorizing biological materials by their physical form and collection method, such as ‘body fluid specimen’ and ‘biopsy specimen’. The sample_type attribute categorizes biological samples based on their neoplastic characteristics, such as ‘Primary Neoplasm’ and ‘Metastatic Neoplasm’ (**Figure 4c**). The 77 unique values under three original attributes were harmonized into 20 standardized terms under two curated attributes. For these two harmonized attributes, completeness actually decreased because values belonging to different/irrelevant attributes had been mixed together in the original data, artificially inflating completeness through the inclusion of irrelevant information. About 61% of the samples have sample_type information, and the majority (52%, n = 59,831) are classified as ‘Primary Neoplasm’, followed by ‘Neoplasm’ (30%, n = 34,998) and ‘Metastatic Neoplasm’ (17%, n = 19,892).

#### Vital Status

Approximately 62% (n = 117,950) of the samples in cBioPortal have vital status information initially spread across twenty different attributes, while there was one attribute (‘OS_STATUS’) with high completeness (59%). This harmonization consolidated 67 original values into three curated values (Dead, Alive, and Unknown), significantly improving consistency, findability, and interoperability.

### *OmicsMLRepoR* for accessing harmonized metadata

Controlled vocabularies (i.e., ontologies) are highly recommended for standardizing metadata descriptions, and several software tools support this implementation(27–30). However, querying data through ontological relationships remains limited because most existing data resources use ontologies primarily for annotation rather than enabling flexible, hierarchy-aware queries. Furthermore, the functionalities of existing tools are often scattered, complex, and not user-friendly. To facilitate the efficient use of our harmonized metadata, we developed the *OmicsMLRepoR*(*31*) Bioconductor package. It provides intuitive functions for querying harmonized metadata through ontological relationships, reshaping data for downstream analysis, and identifying shared ancestor terms.

## Discussion

We address critical challenges in omics data accessibility and reuse through comprehensive, retrospective, and cross-study metadata harmonization. Our approach harmonizes key attributes across nearly 500 studies, which encompass over 210,000 samples from major cancer genomics and metagenomic databases, thereby enhancing their FAIR compliance and AI/ML applicability. Unlike prospective standardization frameworks that impose predetermined schemas, our approach derives data models from an empirical analysis of actual usage patterns across existing datasets. This evidence-based schema design, combined with our flexible attribute schemas (i.e., composite, multi-valued, and *_details attributes), enables the practical integration of metadata while preserving the granular information required for diverse analytical approaches.

A persistent challenge in metadata management is achieving a balance between comprehensiveness and practicality, both during curation and usage. Manual curation often leads to redundancy as curators create new attributes when existing ones do not exactly match or when data schemas become too complex to navigate efficiently. This redundancy scatters related data across multiple attributes, making it difficult for users to locate relevant information. While some data repositories(32) capture metadata in great detail, the resulting complexity makes these resources more difficult to use, requiring a deeper understanding of data models and significant effort for data access and cleaning. Our approach focuses on harmonizing major, high-level attributes, such as demographics, diseases, and treatments, to improve findability while preserving access to all study-specific attributes through their source repositories. This strategy enables researchers to identify larger numbers of similar public datasets while maintaining study-specific details.

The harmonized metadata directly addresses several pressing concerns in omics research. Insufficient representation of minorities in genomic sequencing projects risks widening health disparities, while failure to include sex as a biological variable in AI/ML projects may compromise model validity. Additionally, most cancer omics AI/ML applications typically integrate three to four major omics data types (e.g., genomic, transcriptomic, proteomic, and epigenomic) (33,34), while the incorporation of clinical metadata remains limited (6,33). Our standardized, machine-readable reporting of clinical metadata helps address these critical gaps.

OmicsMLRepo addresses the practical challenge of harmonizing already-collected research data, distinguishing it from prospective standardization frameworks such as OMOP CDM or HL7 FHIR, which require adoption during data collection. While OMOP excels in clinical data standardization and HL7 FHIR enables real-time health data exchange, neither framework is optimized for the retrospective curation of diverse omics research metadata across independent studies. Our approach complements these established frameworks by providing a lightweight, ontology-driven solution specifically designed for already existing data. Especially, the flexibility of our approach can facilitate the integration of heterogeneous data and enable a quick adaptation to evolving standards as new measurement techniques or data collection practices emerge.

While retrospective harmonization addresses immediate needs for existing data, we acknowledge several limitations compared to prospective standardization approaches. Our manual curation process requires substantial resources and introduces a temporal lag between data publication and the availability of the harmonized versions. Prospective adoption of standards during data collection would eliminate these inefficiencies and ensure comprehensive metadata capture from the outset. However, the reality is that vast amounts of valuable research data already exist in non-standardized formats, and achieving consensus and adoption of prospective standards across the diverse omics research community remains challenging. The heterogeneity of research contexts makes it difficult to implement one-size-fits-all prospective standards. Thus, while limitations exist, our approach provides immediate value for existing data while the community works toward better prospective practices. The ideal scenario combines both approaches: using frameworks like OmicsMLRepo to maximize the utility of existing data while simultaneously encouraging the adoption of the OmicsMLRepo data schema as prospective standards for new data collection. Also, to expand this project, it will be critical to develop an automated metadata harmonization pipeline. With recent advances in large language models(34,35), unstructured data may be incorporated into initial metadata curation to further reduce manual effort.

We envision OmicsMLRepo as a pan-repository index system, enabling researchers to browse and identify relevant omics data through metadata without prior knowledge of their existence. The presented work demonstrates a significant advancement and potential for further improvement in making public omics data more accessible and AI/ML-ready while establishing a framework for better metadata management practices in future studies.

## Data Availability

The *OmicsMLRepoR* package is distributed through Bioconductor (https://www.bioconductor.org/packages/release/bioc/html/OmicsMLRepoR.html). The harmonized cMD and cBioPortal metadata tables are available in Zenodo (https://zenodo.org/record/14110745) and can be accessed directly through the *OmicsMLRepoR* package. The data merging schemas and data dictionaries are provided in the Supplementary Tables.

## Author Contributions

Study conception and design: L.W., S.D., S.O.; Data collection and processing: K.L., K.G., S.O.; Pipeline development: K.L., K.G., S.D., S.O.; Analysis and visualization: S.O.; Writing - original draft: S.O.; Writing - review and editing: K.L., K.G., L.W., S.D., S.O.; Funding: L.W., S.D., S.O. All authors approved the final datasets and manuscript.

