## Supplementary Note for "Large-scale Manual Curation and Harmonization of Metadata from Metagenomic and Cancer Genomic Repositories: Challenges and Solutions"

### **Supplementary Note: Practical Guidelines for Retrospective Metadata Harmonization**

These guidelines synthesize lessons learned from harmonizing 468 studies comprising over 210,000 samples across two major omics repositories. While our framework is described in the main text, the following recommendations address practical implementation challenges that harmonization practitioners will encounter when applying similar approaches to their own datasets. These guidelines complement the core methodological innovations presented in the main text by providing detailed best practices for quality control, standardization, and documentation throughout the harmonization process.

**1. Investigate potential reasons for discordance.** It is crucial to distinguish between systematic and random errors, as the cause of discordance determines the appropriate correction strategy. Systematic errors suggest that you should contact data providers to correct the source, preventing future issues. Also, systematic errors should be fixed before applying the majority-rule-based correction. For example, during our harmonization of sex attribute in *cBioPortalData*, we found that 303 out of 337 discordances were from a single study (*luad_msk_npjpo_2021*) and confirmed that it was a mistake at the original data recording step.

**2. Check logical consistency** through cross-sample validation and correction. This process identifies a logical error that needs correction based on established medical knowledge. For example, ‘prostate cancer’ and ‘female’ are impossible combinations for a single sample, because females do not have a prostate and cannot develop prostate cancer.

**3. Resolving granularity mismatches**. We keep the most specific information while offering widely used, higher-level information separately as needed. Data from different sources often contains information at varying levels of detail; some may use broad categories while others employ highly specific terms. Ontologies help address this granularity mismatch by providing hierarchical structures that allow you to identify an appropriate level of specificity for harmonization. We accomplished this dual-level approach by creating both standard/main attributes and their corresponding 'details' versions. This eliminates the traditional trade-off between preserving granular detail and achieving broad compatibility, ensuring that no meaningful information is discarded during the harmonization process. However, implementing this strategy requires domain knowledge, as terms can often appear in multiple ontological branches, potentially leading to multiple valid ancestor terms. These must be carefully evaluated to select the most appropriate hierarchical path for preserving the original meaning and facilitating harmonization purposes.

**4. Maintain a consistent naming and formatting schema**, which is essential for scalability, interpretability, and automated processing. First, we suggest using consistent prefixes or suffixes in file and attribute names. Additionally, we recommend making both file and attribute names human- and machine-readable; i.e., avoid using programming language keywords, special characters, and spaces. Consistent value formatting examples include using a single standard NA (not a mix of NULL, missing, -, or 999) and consistently formatting boolean values (e.g., yes/no or true/false).

**5. Quality metrics and validation.** We used the four criteria to assess the improved quality of metadata from semi-structured data (**Figure 1b**). However, the interpretation of these quality improvements is context-dependent and may vary depending on the characteristics of your input data. For example, increasing standardization might initially reduce completeness when strict validation rules exclude included-but-non-conformant values. Similarly, improving consistency may reduce the number of unique values as detailed free-text entries are mapped to standardized categorical terms. Different domains of data may require distinct evaluation metrics, such as reference range alignment (ensuring clinically meaningful normal/abnormal flags) or a scope of allowed values. While there is no universal, predefined rule for metric selection, having proper quality metrics is essential for evaluating the harmonization strategy, detecting errors, and iteratively refining the harmonization process to achieve an optimal balance across all quality dimensions while maintaining usability.

**6. Data provenance tracking and documentation are helpful for both you and others.** Comprehensive documentation captures not just what was done, but why specific decisions were made during the harmonization process. This contextual information becomes invaluable when original team members leave, when revisiting old projects, or when training new researchers on established protocols. Detailed tracking also enables tracing back through the process when discrepancies or unexpected results emerge. This audit trail enables the identification of exactly where issues occurred and facilitates targeted corrections without requiring the restart of the entire harmonization process.
